# A 3-dimensional molecular cartography of human cerebral organoids revealed by double-barcoded spatial transcriptomics

**DOI:** 10.1101/2023.01.18.524520

**Authors:** Gwendoline Lozachmeur, Aude Bramoulle, Antoine Aubert, François Stüder, Julien Moehlin, Lucie Madrange, Frank Yates, Jean-Philippe Deslys, Marco Antonio Mendoza-Parra

**Affiliations:** UMR 8030 Génomique Métabolique, Genoscope, Institut François Jacob, CEA, CNRS, University of Evry-val-d’Essonne, University Paris-Saclay, 91057 Évry, France; Service d’Etude des Prions et des Infections Atypiques (SEPIA), Institut François Jacob, Commissariat à l’Energie Atomique et aux Energies Alternatives (CEA), Université Paris Saclay, Fontenay-aux-Roses, France; Sup’Biotech Engineering School - CellTechs Team, Villejuif, France

**Keywords:** spatial transcriptomics, cerebral organoids, gene regulatory programs, DNA arrays

## Abstract

Spatially-resolved transcriptomics is revolutionizing our understanding of complex tissues, but their current use for the exploration of a few sections is not representative of their 3-dimensional architecture. In this work we present a low-cost strategy for manufacturing molecularly double-barcoded DNA arrays, enabling large-scale spatially-resolved transcriptomics studies. We applied this technique to spatially resolve gene expression in several human brain organoids, including the reconstruction of a 3-dimensional view from multiple consecutive sections, revealing gene expression divergencies throughout the tissue.

## Main text

Understanding tissue complexity by the characterization of gene programs associated with the various cells/cell types that are composing them represent a major challenge in systems biology. Recent developments in spatially-resolved transcriptomics (SrT) are revolutionizing the way to scrutinize tissue complexity without losing its spatial architecture. Among the various available strategies, those based on the use of a “barcoded” physical support (for capturing the molecular information), combined with the use of next-generation DNA sequencing and bioinformatics demultiplexing processing (for tracing-back the local positioning of the assessed transcriptomes) are providing means to landscape large tissue sections and interrogate gene expression in an unbiased manner ^1^.

Barcoded DNA microarrays, initially described by Stähl et al for SrT applications ^2^, allow to capture messenger RNA thanks to DNA probes presenting a poly(T) sequence, and provide local transcriptome positional information thanks to a unique molecular barcode associated with each of the DNA probes. While powerful, this approach is rather expensive, notably due to the high number of unique DNA probes required for their manufacturing.

Herein we present an improved strategy for manufacturing molecularly barcoded DNA arrays for SrT. Instead of printing long DNA oligo-nucleotides (>100 nts) presenting a unique 18-mer barcode sequence and a 20nt-length poly(T) capture region as described by Stäh et al ^2^, we have engineered a strategy based on the use of two DNA oligonucleotides harboring two distinct sets of molecular barcodes (BCri and BCcj; **Figure 1A**). In addition, one of them hosts an amino C6 linker modification at the 5’-end for UV-crosslinking, a T7 promoter sequence, and a 30 nt length adapter at the 3’-end, previously used for Gibson assembly reactions ^3^. A complementary Gibson sequence is retrieved at the 3’-end of the second printed DNA oligonucleotide, as well as 20nt-length poly(A) extremity at its 5’end. During the manufacturing process, the first types of oligonucleotides are printed as rows, such that each row presents oligonucleotides with a different barcode sequence (BCr1, BCr2,…BCri; **Figure 1A**). Then, the second types of oligonucleotides are printed as columns, on top of the previously printed oligonucleotides, such that each column contains oligonucleotides with a different barcode sequence (BCc1, BCc2,…BCcj).

**Figure 1.**
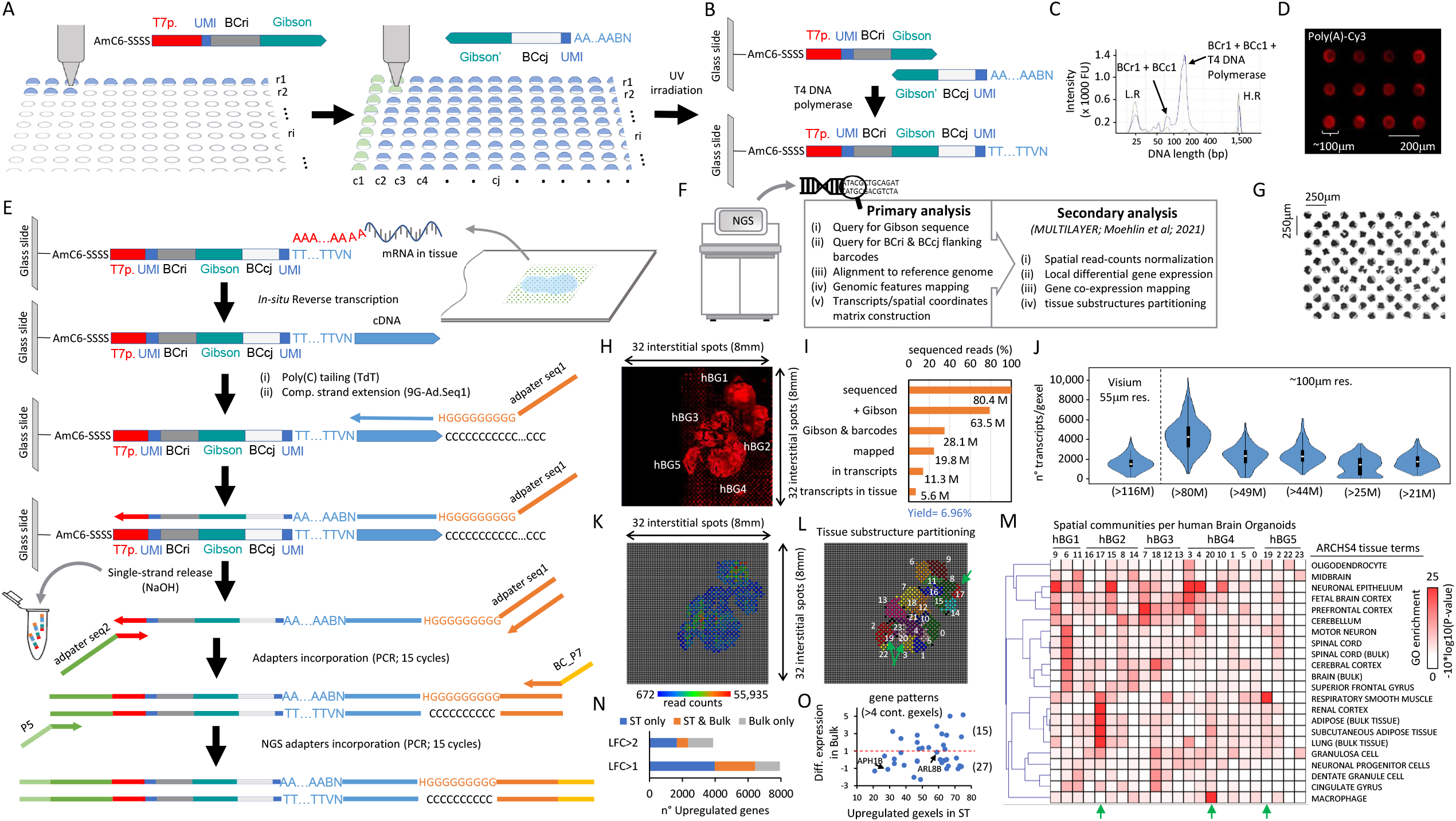
Double-barcoded DNA arrays manufactured for spatially-resolved transcriptomics (SrT). **(A)** Strategy for printing DNA arrays based on the deposition of two types of oligonucleotides harboring distinct molecular barcodes (per rows: BCri; per columns: BCci). Both oligonucleotides share a complementary sequence (Gibson). The 5’-end of the BCri oligonucleotide contains an amino C6 linker modification, followed by 4 “S” (Cytocine or Guanine) nucleotides, for UV-crosslinking. **(B)** Once BCcj primers are deposited on top of the printed BCri oligonucleotides, the glass slide is UV irradiated for covalent crosslinking, followed by probes’ elongation thanks to their complementary Gibson region. This elongation process (T4 DNA polymerase) generates a long probe composed by a T7 promoter (T7p), two unique molecular identifiers (UMI), two molecular barcodes flanking the complementary Gibson sequence and a poly(T). **(C)** Electropherogram demonstrating the generation of a long probe (116nt length) after T4 DNA Polymerase elongation. **(D)** DNA array scan (TRITC filter) after incubation of a double-barcoded DNA array with poly(A)-Cy3 labelled molecules. **(E)** NGS library preparation strategy for capturing messanger RNA (mRNA) from tissues deposited on top of the double-barcoded manufactured DNA arrays. **(F)** Bioinformatics pipeline in use for spatially resolved transcriptomics. **(G)** Micrograph illustrating a part of a manufactured DNA array composed by 32×32 printed spots in an interstitial manner, leading to a density of 2048 different probes. **(H)** Scan of a DNA array (TRITC filter) hosting a cryosection of multiple human brain organoids (hBGs embedded together), after cDNA labeling with dCTP-Cy3. **(I)** Fraction of sequenced reads retrieved from library preparation in (H) after the primary bioinformatics analysis. Yield corresponds to the ratio between the reads retrieved in transcripts in the tissue region (5.6 Million) and the initial number of sequenced reads (80.4 Million). **(J)** Violin plots illustrating the number of transcripts per gexel (gene expression elements, i.e. a spatial spot) retrieved in several SrT assays performed with double-barcoded manufactured DNA arrays at different sequencing depths (80 to 20 million reads), in comparison with an assay with commercial DNA array (Visium 10X Genomics). Note that the Visium assay has been performed with >100 million reads. **(K)** Digitized view of the SrT assay performed in (H) generated with MULTILAYER from the 5.6 million reads retrieved in known transcripts in tissue regions. **(L)** Molecular tissue substructure partitioning performed by MULTILAYER based on gene co-expression pattern signatures (≥ 4 contiguous gexels per pattern). Each color corresponds to one of the 24 revealed substructures. **(M)** Tissue terms enrichment analysis per substructures retrieved in (L) and classified by their localization within the identified hBGs. Green arrows: Spatial substructures (19, 20, 17) enriched for tissue terms not related to neuro-ectodermal differentiation. Spatial locations of these structures are also highlighted in (L) (green arrows). **(N)** Comparison between Spatial (ST; in H) and bulk transcriptomics assessed in hBGs. LFC: Fold-change in log2. **(O)** Comparison between the number of upregulated gexels in ST retrieved in gene patterns (>4 contiguous gexels) and the differential expression detected in bulk transcriptomics. Red dashed line highlights a LFC = 1. Two genes (APH1B and ARL8B) are highlighted as not found differentially expressed in bulk but presenting an important number of upregulated gexels in the ST assay.

After UV irradiation, printed DNA arrays are covered with a solution containing T4 DNA polymerase for elongating the hybridized oligonucleotides, giving rise to 116 nt length DNA probe presenting a poly(T) sequence at the 3’-end and two distinct barcodes providing positional information (row/column coordinates) (**Figure 1B&C**). The capacity of the double-barcoded DNA arrays to capture poly(A) sequences was verified by their exposure to Poly(A)-Cyanine-3 (Cy3) labeled oligonucleotides (**Figure 1D**). Importantly, the aforementioned manufacturing strategy of double-barcoded DNA arrays requires only 64 oligonucleotides for generating 1024 different DNA probes; or only 128 oligonucleotides for reaching a density of 4096 different probes, similar to the currently improved DNA arrays commercialized by 10xGenomics. Hence, the production costs of the double-barcoded arrays are 16 to 32-fold lower than those presenting a single 18-mer molecular barcode.

In order to use the aforementioned double-barcoded DNA arrays for spatially-resolved transcriptomics, we have engineered a molecular biology strategy in which, the messenger RNA retrieved in tissue sections is converted in complementary DNA (cDNA) in-situ, followed by a poly(C) tailing with Terminal transferase, as described in previous NGS-library preparation strategies ^4,5^ (**Figure 1E**). The poly(C) sequence is used for hybridizing a poly(G)-oligonucleotide presenting a known adapter at its 5’-end (Adapter seq1), and elongated for generating a complementary DNA strand harboring the captured cDNA sequence, as well as the double-barcoded information. Finally, the complementary strand is released by alkaline treatment and recovered from the DNA array for its transfer into an Eppendorf tube. A second adapter attached to a T7 promoter sequence is used together with a complementary sequence to the Adapter seq1 for a first PCR DNA amplification step, followed by a second amplification process incorporating the required adapters for Illumina NGS sequencing (P5; P7 sequences; **Figure 1E**).

In addition to the molecular biology strategy for SrT assays, we have generated a bioinformatics pipeline targeting the retrieval of the Gibson sequence flanked by two molecular barcodes prior to the alignment to the reference genome, mapping of genomic features, and the reconstruction of a spatial coordinates matrix associated to the retrieved read-counts per transcripts (**Figure 1F**). The outcome of this primary analysis is processed by our previously described tool MULTILAYER, able to normalize the spatial read-count levels, identify differentially expressed genes in a local context, detect gene co-expression patterns and perform molecular tissue substructure partitioning ^6,7^.

The double-barcoding strategy has been used for manufacturing DNA arrays composed of 2048 unique DNA probes (32×32 spots printed twice in an interstitial manner; **Figure 1G**). Then, multiple human brain organoids (hBG; 4 months of culture) were cryo-sectioned together and deposited on top of the manufactured DNA arrays. In-situ reverse transcription was revealed by the incorporation of dCTP-Cy3 (**Figure 1H**). Next-generation sequencing of the corresponding SrT library revealed that from a total of ∼80 million reads, 63 million presented the Gibson sequence, but only ∼28 million presented in addition the required flanking positional barcodes (**Figure 1I**). Furthermore, ∼20 million reads were mapped to the human genome, from which 5.6 million matched with known transcripts and were retrieved within tissue regions, leading to a yield of ∼7% relative to the total mapped reads. The analysis of multiple other sections at various sequencing-depth levels revealed a yield of spatially captured transcripts ranging between 3.5 to 7%, equivalent to that obtained when using commercial DNA arrays (Visium 10x Genomics (**Supp. Fig. S1**)). Importantly, our double-barcoded arrays presented ∼4000 transcripts per gexel (local gene expression pixels) in average for a sequencing depth of ∼80 million reads; while the assay with Visium DNA arrays revealed less than 2000 transcripts per gexel, in despite of the use of >116 million reads. This can be explained by the difference in the number of printed spots and resolution (2048 spots in our double-barcoded arrays and ∼5000 spots for Visium arrays) (**Figure 1J**). Indeed, the use of >40 million reads for SrT with the double-barcoded arrays allowed us to obtain >2000 transcripts/gexel (**Figure 1J & Supp. Fig. S1**).

A digitized view of the analyzed hBGs revealed a SrT map presenting raw read counts ranging between 672 and >55 thousand per gexel (**Figure 1K**). Tissue partitioning, based on localized gene co-expression patterns (≥ 4 contiguous gexels per pattern; Tanimoto Similarity threshold: 25%), revealed 24 molecular tissue substructures (**Figure 1L and Supplementary Figure S2**). Tissue terms enrichment analysis applied to co-expressed genes retrieved in partitioned regions revealed signatures associated with terms like neuronal epithelium or fetal brain cortex as being enriched in most of the hBGs, while others like oligodendrocyte, Spinal cord, or neuronal progenitor cells being more restricted to a given hBG (**Figure 1M**). Such differences strongly support the well-described heterogeneity within hBG cultures, further supported by the observation of some tissue substructures presenting tissue terms devoid of neuroectoderm-derived signatures (green arrows; 17, 19, 20; **Figure 1L&M**). Finally, a direct comparison of the up-regulated genes detected with the double-barcoded SrT strategy with a bulk transcriptome mapping revealed ∼30% of overlap (**Figure 1N**). Such difference can be explained by the inherent heterogeneity within hBGs, as well as the sensitivity of SrT assays to reveal local signatures which are averaged in bulk assays. Indeed, Among the 42 up-regulated gene expression patterns detected within the analyzed hBGs, 27 are not found up-regulated in bulk (**Figure 1O**). Among them, genes like APH1B (coding for a gamma-secretase subunit, and recently considered as a biomarker for Alzheimer’s disease ^8^) or ARL8B (GTPase playing a major role in the positioning of interstitial axon branches ^9^) presented >30 upregulated gexels in the SrT map but were not found induced in the bulk transcriptome assay (**Figure 1O & Supplementary Figure S3**).

One of the major interests to count with a low-cost production of DNA arrays for SrT is to be able to perform a large number of assays, for instance for reconstructing a 3-dimensional view of the tissue of interest. Herein, we analyzed nine consecutive sections of two human brain organoids (9 Months of culture), covering 540 microns of the tissue thickness (**Figure 2A**), deposited on double-barcoded DNA arrays (2048 probes; **Figure 2B & Supplementary Figure S4**). Their processing with the help of MULTILAYER allowed identifying several differentially expressed genes across sections, among them CNTNAP2, a member of the neurexin family, involved in synapse formation and essential for neuronal development ^10^ (**Figure 2C**). In addition to CNTNAP2 which appeared locally over-expressed across all sections (**Figure 2D**), we have found other 277 genes presenting the same behavior (**Figure 2E**). Furthermore, between 60 to 500 genes were upregulated specifically in each section, while most of them (up to 2500 genes) appeared upregulated in more than one section (**Figure 2E**). Interestingly, these “shared” genes, appeared upregulated preferentially up to three consecutive sections across the tissue, covering a distance of ∼200 microns, corresponding to ∼10 cells on average (**Figure 2F**).

**Figure 2.**
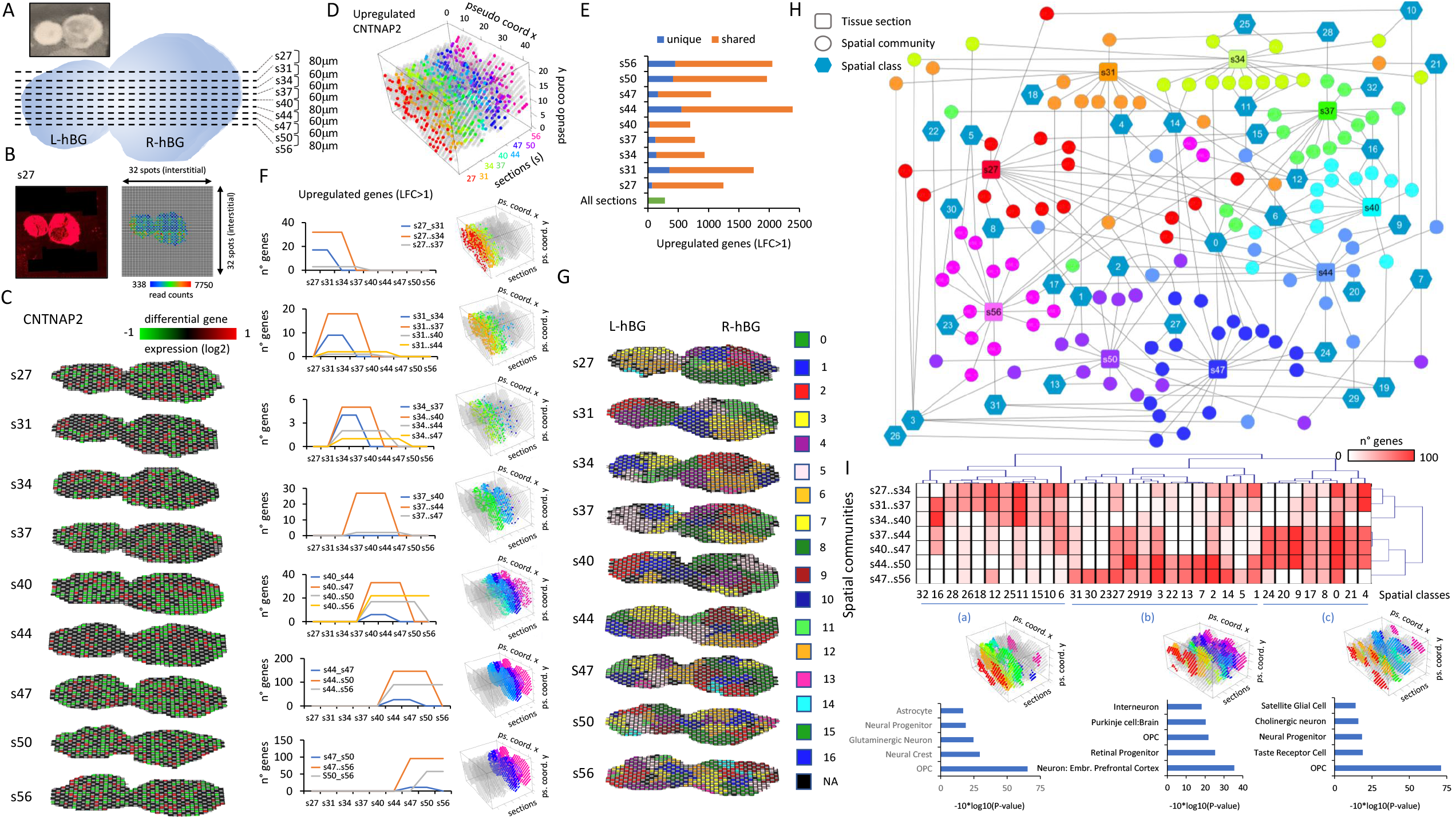
3-Dimension spatial transcriptomics view of human brain organoids. **(A)** Top: micrograph displaying a tissue section of two human brain organoids (hBG) embedded together. Bottom: Scheme of the consecutive sections (s27 till s56) collected across both hBGs (Left: L-hBG; right: R-hBG). **(B)** Left: Scan of a DNA array (TRITC filter) hosting a cryosection (section 27) after cDNA labeling with dCTP-Cy3. Right: SrT digitized view of the corresponding section displaying the number of reads counts per gexel. **(C)** Differential expression view of the gene CNTNAP2 across all 9 sections, as revealed by MULTILAYER. 3-dimensional view of the upregulation of CNTNAP2 across all sections (Log2 fold-change (LFC)>1). Pseudo-coordinates are computed by aligning all sections relative to the center of the L-hBG and R-hBG. Colored gexels highlight CNTNAP2 upregulation per tissue section. **(E)** Number of upregulated genes (LFC>1) across all sections (green bar-plot), specific to a given section (blue bar-plots) or shared among at least two sections (orange bar-plots). **(F)** Left: Upregulated genes in consecutive sections. Right: 3-dimensional view of the upregulated genes across sections. Colored sections coincide with the color-code per section displayed in (D). **(G)** Molecular tissue stratification based on gene co-expression pattern signatures performed by MULTILAYER (≥ 3 contiguous gexels per pattern; Tanimoto Similarity threshold: 10%) A total of 16 different spatial communities (subregions) were identified across all sections. **(H)** Network representation of the various spatial communities detected per tissue section and their association to spatial classes, defined as spatial communities sharing common gene co-expression patterns. Colored sections coincide with the color-code per section displayed in (D). **(I)** Top panel: Clustering of the number of genes per spatial classes and associated to 3 consecutive sections revealing three major groups (“a”, “b” & “c”) (Pearson correlation). Middle panel: 3-dimensional view of the spatial classes per tissue section retrieved in all three groups. Low panel: Cell-type enrichment analysis associated with all three groups (Panglao and CellMarkerAugmented DBs).

Partitioning consecutive sections in molecular substructures based on gene co-expression patterns (≥ 3 contiguous gexels per pattern; Tanimoto Similarity threshold: 10%) revealed 16 spatially localized communities (**Figure 2G**). Their comparison across sections has been performed from their gene co-expression content per spatial community, leading to the identification of 33 spatial classes. A network representation of these relationships revealed the presence of spatial classes shared among multiple tissues (e.g. class “0”), others being specific to consecutive sections (e.g. class “16”) (**Figure 2H**). Clustering upregulated genes retrieved in three consecutive sections and associated with the aforementioned spatial classes revealed the presence of three major groups based on their spatial localization (**Figure 2I**). These groups are enriched for a variety of neuronal subtypes (Glutamatergic: Group “a”; Cholinergic: Group “c”), including specialized terms like Retinal progenitors (group “b”) or Taste receptor cells (group “c”), but also glial cells (Astrocytes: Group “a”; Oligodendrocyte precursors (OPC): Group “a”, “b” and “c”) in agreement with a complex tissue architecture due to the variety of cell types but also to their spatial localization (**Figure 2I & Supplementary Figure S5**).

Overall, we have provided herein a 3-dimensional view of the molecular complexity retrieved within human cerebral organoids. As far as we know, together with the work performed by Ortiz et al (revealing a molecular atlas of the adult mouse brain ^11^), this is the second report focused in the integration of multiple consecutive spatial transcriptomics; most likely due to the elevated costs associated to these type of assays. We anticipate that the manufacturing strategy for producing double-barcoded DNA arrays reported herein will provide means to democratize the use of SrT assays, for instance for reconstructing 3-dimensional molecular landscapes of complex tissues.

## Methods

### Cerebral Organoid formation

Human induced Pluripotent Stem Cells were cultured in mTeSR Plus Medium (100-0276 Stem Cell) + Penicillin/Streptomycin (P/S) 1% (15140122 Thermo Fisher Scientific) on Matrigel coating (354277 Corning). Cerebral organoids were cultured by forming embryoid bodies (EBs) using hanging drops (20,000 cells per drop of 22 µL) in **EB Formation Medium** (for 50 mL: 40 mL DMEM/F12 (11330-032 Thermo Fisher Scientific), 10 mL KOSR (10828-028 Thermo Fisher Scientific), 500 µL GlutaMAX (35050-038 Thermo Fisher Scientific), 500 µL MEM-NEAA (11140-050 Thermo Fisher Scientific), 500 µL P/S supplemented just before use with bFGF (PHG0264 Thermo Fisher Scientific) at 4 ng/mL, Y-27632 dihydrochloride (ROCK Inhibitor, 72304 Stem Cell Technologies) at 50 µM, SB-431542 (72232 Stem Cell Technologies) at 10 µM and LDN-193189 (72147 Stem Cell Technologies) at 100 nM). After 2-3 days as hanging drops, EBs were transferred in 24 well Ultra Low Attachment plates (Thermo Fisher Scientific P24; ref. 144530 “ untreated Nunc”) with **EB Formation Medium** (same as above except that ROCK Inhibitor is used at 20 µM), to refresh 2 days after transfer. After 1 week in **EB Formation Medium**, EBs were transferred in **Neural Induction Medium** (for 50 mL: 48.5mL DMEM/F12, 500 µL N2 Supplement (17505-048 Thermo Fisher Scientific), 500 µL GlutaMAX, 500 µL P/S, 500 µL MEM-NEAA) until embedding in Matrigel® Matrix (354230 Corning) using an Organoid Embedding Sheet (08579 Stem Cell Technologies). After embedding, droplets were put in 6-well Ultra Low Attachment Plates (Thermo Fisher Scientific P6 : REF 150239 “untreated Nunc”) in **Expansion Medium** without vitamin A (for 50 mL: 24 mL DMEM/F12, 24 mL Neurobasal Medium (21103-049 Thermo Fisher Scientific), 500 µL GlutaMAX, 500 µL P/S, 500 µL B27 without Vitamin A Supplement (12587-010 Thermo Fisher Scientific), 250 µL N2 Supplement, 250 µL MEM-NEAA, 12.5 µL Insulin solution, 0.5 µL 2-Mercaptoethanol). After 4 days in **Expansion Medium**, Organoids were transfered to **Maturation Medium with Vitamin A** (same as Expansion Medium, except B27 without Vitamin A is replaced by B27 Supplement (17504-044 Thermo Fisher Scientific)) and cultured under shaking at 60 rpm (Infors Celltron Orbital Shaker) with refresh of medium once a week. Human brain organoids used in this study were collected either at 4 or 9 months of culture by snap-freezing in isopentane and embedded in OCT prior cryosectioning.

### Bulk transcriptomics for human brain organoids

For Bulk RNA-seq transcriptomics, a human brain organoid cultured during 4 months has been dissociated in TRIzol RNA isolation reagent (Thermo Fisher Scientific ref: 15596026). Extracted RNA has been processed with the NEBNext Ultra II RNA Library Prep Kit for Illumina (E7770). Human induced pluripotent stem (iPS) cells were also processed with TRIzol RNA isolation reagent and NEBNext Ultra II RNA Library Prep Kit. Libraries were sequenced within the French National Sequencing Center, Genoscope (150-nt pair-end sequencing; NovaSeq Illumina).

Both bulk human brain organoid and iPS control sequenced samples were aligned to the human reference genome (hg19; Bowtie 2.1.0 under default parameters). Mapped reads were associated with known transcripts with featureCounts ^12^. Differential expression analysis between bulk human brain organoid and iPS control readouts was done with DESeq2 R package.

### DNA arrays manufacturing

DNA arrays for spatially resolved transcriptomics were manufactured by depositing two types of complementary oligonucleotides, herein defined as “Barcode for rows (BCr)” and “Barcode for columns (BCc)” hosting sequences. The BCr oligonucleotide presents an amino C6 linker at the 5’ extremity, followed by four G or C nucleotides (S), a T7 promoter sequence (GACTCGTAATACGACTCACTATAGGG), a unique molecular idenfitier (UMI: WSNNWSNNV), a molecular barcode (8 nucleotides) associated to the printed row in the DNA array, and a 30 nucleotides long adapter sequence with a GC-content of 40%, previously used for Gibson assembly reactions ^3^ (herein named as Gibson sequence: “ACATTGAAGAACCTG-TAGATAACTCGCTGT”).

Similarly, the BCc oligonucleotide is composed by “NBAAAAAAAAAAAAAAAAAAAA” sequence at the 5’ extremity (where B corresponds to any nucleotide except A), a unique molecular idenfitier (UMI: WSNNWSNNV), a molecular barcode (8 nucleotides) associated to the printed column in the DNA array, and a complementary Gibson sequence.

Oligonucleotides are diluted to 5 µM in presence of sciSPOT Oligos B1 solution (Scienion CBD-5421-50) in a 96 wells plate and stored at -20°C for long storage.

DNA arrays were printed onto Superfrost Plus Adhesion Microscope Slides (J7800AMNZ Epredia). For generating DNA arrays composed by 2048 different probes (177 µm pitch distance; ∼100 µm printed spot), 32 BCr oligonucleotides presenting a unique molecular barcode were printed per row, by depositing ∼250 pL (250 µm between contiguous spots) with a DNA pico-litter spotter (Scienion sciflexarrayer S3). Similarly, 32 different BCc oligonucleotides were printed per column, by spotting ∼250 pL of each of them on top of the previoulsy prinded BCr nucleotides. Then, the same 32 BCr oligonucleotides were printed per row with a shifted position of 125 µm in both axis (i.e. interstitial printing), followed by the deposition of other 32 different BCc oligonucleotides printed per column. Hence, we needed in total 32 unique BCr (printed twice) and 64 unique BCc oligonucleotides.

Finally, fiducial borders were printed per DNA array, by adding three row/columns of spots with the used pitch distance (250 µm) by printing a BCr1 oligonucleotide presenting a Cy3-label at the 3’-extremity.

UV irradiation (254 nm; 5 min) has been applied for crosslinking, followed by T4 DNA polymerase elongation (15 µL /printed region with coverslips: 0.03 U/µL T4 DNA Polymerase New England Biolads M0203L; 0.2 mM dNTPs Thermo Fisher Scientific R0141/ R0151 / R0161 / R0171; 37°C; 1hour). To monitor the double-strand DNA elongation performance, a quality control assay is systematically done within a batch production (18 slides per batch; 3 printed regions per glass slide (8×8 mm size; 250 µm pitch distance)) by incorporating a dCTP-Cy3 nucleotide during the elongation (dATP/GTP/TTP at 52 µM, dCTP at 50 µM and dCTP-Cy3 at 20 µM (Cytivia PA53021)), and analyzed by fluorescent microscopy. After elongation, slides have been washed with 0.1X SSC buffer (Sigma S6639), and ddH_2_O, and finally spin dried (Labnet slide spinner C1303-T-230V, 4800 rpm). DNA array slides have been stored at +4°C in a sealed container.

### Organoids tissue cryosectioning and in-situ reverse transcription

Tissue samples were cryosectionned (20 µm sections collected at -20°C) and deposited on the DNA-arrays defined regions. During this process, slides were kept inside the cryostat chamber in order to conserve the integrity of the RNA and at -80°C for long storage.

Tissue sections were fixed with 4% Para-formaldheyde (PFA) (Life technologies 28908) in 1X PBS (Life technologies 70011-036) at room temperature during 10 minutes, then washed twice with 1X PBS, followed by a wash with double-distilled water (ddH_2_O).

Tissues deposited on DNA arrays were covered with ddH_2_O and heat at 65°C during 5 minutes, followed by a fast chilling procedure (−20°C; 2 minutes). Remaining water was replaced by hybridization mix (10 U RNAse Inhibitor(New England Biolads M0314) in 6X SSC), and incubated at room temperature for 30 minutes. DNA arrays were washed by dipping them into a 50 mL falcon of 0.1X SSC then into a 50 mL falcon of ddH_2_O. Finally, tissue sections were air dried during 10 minutes.

To avoid evaporation, reverse transcription (RT) is realised within sealed chambers (Thermoscientific small size; AB-0576). DNA arrays presenting tissue sections were incubated with RT mix (1 U/µL of SuperScript IV at (18090200 Life Technologies), 1X SSIV buffer (18090200 Life Technologies), 5 mM DTT (18090200 Life Technologies), 20 µg/mL actinomycin D (A1410-2MG Sigma), 0.2 mg/mL Bovine Serum Albumine (BSA) (Sigma Merk A9418-10G), a mix of dATP/dTTP/dGTP (130 µM each), 125 µM of dCTP and 5 µM dCTP-Cy3, 1 U/µL of RNAse Inhibitors) at 42°C overnight with a heated lid at 70°C. Next day, slides were washed by dipping them into a 50 mL falcon of 0.1X SSC, then into a 50 mL falcon of ddH_2_O and spin-dried. Finally, DNA arrays were scanned with the TRICT filter to reveal the presence of the fiducial borders as well as the physical position of tissue sections on top of the printed DNA arrays and revealed by the newly synthesized cDNA.

After imaging, tissue sections were digested during 1 hour at 37°C and at 300rpm (20 mg/mL proteinase K (Invitrogen 4333793) at 7.5 mg/mL in PKD buffer (Quiagen 1034963)). Slides were washed in containers containing large volumes and under agitation (400 rpm) as following: 100 mL of preheated buffer 1 (2X SSC and 0.1X SDS (Sigma 71736); 50°C) during 10 minutes, 5 minutes with buffer 2 (0.2X SSC; room temperature), 5 minutres with buffer 3 (0.1X SSC; room temperature). Finally the slides were washed by dipping them in a 50 mL falcon containing ddH_2_O and spin-dried. DNA arrays were inspected at this stage to make sure that tissues was completely removed. DNA arrays were incubated with a freshly prepared 0.1 N NaOH solution during 10 minutes at room temperature, washed with the same solution, then neutralized for 2 minutes (for 100µL of 0.1 N NaOH, add 11.8 µL of 10X TE and, 6.5 µL of 1.25 M acetic acid). Finally, slides were washed by dipping them into a 50 mL falcon of 0.1X SSC, then into a 50 mL falcon of ddH_2_O and spin-dried.

### cDNA’s complementary strand synthesis

DNA arrays were incubated with a poly-C tailing mix (0.6 U/µL TDT (New England Biolads M0315), 1X terminal transferase buffer (New England Biolads BO315), 0.3X CoCl_2_ (New England Biolads B0252), 0.2 mM dCTP) during 35 minutes at 37°C, then 20 minutes at 70°C to deactivate the enzyme and cooled at 12°C. Under these conditions, the length of the poly-C tail should be of ∼20 nts in average ^4^. Then, slides were washed in a 50 mL falcon containing 0.1X SSC, then in a falcon containing ddH_2_O prior to be spin-dried.

To generate a complementary DNA strand, DNA arrays were incubated with a 2^nd^ strand sythesis mix (0.1 U/µL klenow exo-(Biolads M0212), 0.2 mg/mL BSA, 1X NEB2 buffer (New Englad Biolads B7002), 0.5 mM dNTPs) in presence of an oligonucleotide presenting a complementary sequence for the poly-C tailed sequence, as well as a 5’-extremity providing an adapter sequence (1µM conc: “GTTCAGACGTGTGCTCTTCCGATCTGGGGGGGGGH”). Second strand synthesis was performed within sealed chambers at 47°C during 5 minutes (primer anhealing), 37°C during 1 hour (extension), 10 minutes at 70°C (enzyme deactivation) and cooled et 12°C. Slides were washed in a 50 mL falcon containing 0.1X SSC and a second falcon containing ddH_2_O.

Newly synthesized complementary DNA strands were recovered by incubating DNA arrays with 100 µL of 0.1N NaOH solution. After 10 minutes of incubation, NaOH solution has been collected in a 1.5 mL eppendorf, DNA arrays were washed with other 100 µL of 0.1N NaOH solution and collected within the same eppendorf. The 200 µL collected NaOH solution has been neutralized with 23.6 µL of 10X TE and 13 µL of 1.25 mM acetic acid. DNA arrays were also neutralized with 100µL 0.1N NaOH solution mixed with 11.8µL 10X TE and 6.5µL 1.25mM acetic acid, then washed with 0.1X SCC and ddH_2_O for potential reuse if required.

Collected solution has been ethanol precipitated (2X EtOH volume, 200 mM NaCl, 1µL of 35 mg/mL glycogen; 1 hour at -20°C; centrifuged at 12000 rpm, washed with 70% EtOH) and resuspended in 23 µL of ddH_2_O for 15 minutes at room temperature.

### cDNA capture validation by quantitative PCR and illumina sequencing library preparation

The 23 µL of cDNA’s complementary-probe strand solution in ddH_2_O has been mixed with 25 µL of Q5 hot start high fidelity 2X master mix(M0464L), 1 µL adapter seq1 primer (GTTCAGACGTGTGCTCTTCCGATCT; 0.02 µM final conc.) and 1 µL adapter seq2 primer containing a part of the T7-promoter sequence (TACACTCTTTCCCTACACGACGCTCTTCCGATCTGACTCGTAATAC; 0.02 µM final conc.).

This mix has been PCR amplifyied (98°C, 30 sec; 15 cycles: 98°C 10 sec; 65°C 75 sec; then 65°C 5 minutes; 12°C hold), cleanned at 0.9X ratio with SPRIselect beads (Beckman Coulter B23318) and resuspended in 24µL of ddH_2_O.

A second round of PCR amplification has been performed by mixing the 24 µL of adaptors linked cDNA’s complementary-probe strand with 25 µL 2X Q5 high fidelity ot start master mix, 0.5 µL of universal primer and 0.5 µL of index primer (each at 0.5µM concentration; NEBNext multiplex oligos for Illumina index primers set 1 E7335). This mix has been amplified for 15 cycles (same amplification conditions as before), then cleanned at an 0.8X ratio with SPRISelect Beads (this ratio is used to get rid off fragments < 200nt because the

DNA-array probes added to the illumina sequencing adaptors and barcodes is 223nt long without capturing any RNA). Final elution has been done in 25 µL of ddH_2_O.

5 µL of this library are used for quantitative PCR evaluation, targeting known transcript sequences (poly-A proximal regions), and the remaining 20 µL of this library is used for Illumina sequencing (150 nts paired-ends sequencing; NovaSeq).

### Bioinformatics processing

Primary analysis has been performed with our in-house developed tool SysISTD (SysFate Illumina Spatial transcriptomics Demultiplexer: https://github.com/SysFate/SysISTD).

SysISTD takes as entry paired-end sequenced reads (fastq or fastq.gz format), and two TSV files, the first one containing the sequence of the molecular barcodes associated to the rows or column in the printed arrays and the second file presenting the physical position architecture of the spatial barcodes. SysISTD search for the Gibson sequence (regex query), then for two neighboring barcodes. Paired-reads presenting these features were aligned to the human genome (hg19) with Bowtie2, and known transcripts were annotated with featureCounts ^12^.

As outcome, SysISTD generated a matrix presenting read counts associated to physical coordinates in the array in columns and known transcripts ID in rows. To focus the downstream analysis to the physical positions corresponding to the analyzed tissue, we used an in-house R script taking as entry an image of the DNA array scanned with the TRICT filter after the reverse transcription step. Indeed, this image reveals the presence of the fiducial borders and the cDNA within the tissue. Specifically, we upload to R a croped image within the fiducials (imager package) and we use the “px.flood” function to retrieve the pixels associated to the tissue. Finally we applied a pixel to gexel coordinates conversion prior to cross this information with the outcome of SysISTD.

The “tissue-focused” matrix presenting read counts associated to spatial coordinates in columns, known transcripts in rows was processed with our previously described tool MULTILAYER ^6,7^. For human brain organoids processed in Figure 1, tissue partitioning has been performed with ≥ 4 contiguous gexels per pattern; Tanimoto Similarity threshold: 25%; while for the 3-dimensional tissue reconstruction in Figure 2, tissue partitioning has been performed with ≥ 3 contiguous gexels per pattern; Tanimoto Similarity threshold: 10%.

To compare contiguous sections, the “batch option” of MULTILAYER has been used, allowing to compute spatial classes in addition to spatial communities. Spatial classes, Spatial communities and tissue relationships were visualized within Cytoscape.

3-dimensional view of upregulated genes across sections were generated with an in-house R script (library “rgl”). Prior visualization, spatial gexels across sections were aligned relative to the center of the left human brain organoid (L-hBG) and the angle between the horizontal axis and a vector between the centers of the L-hBG and the R-hBG. Pseudo coordinates were computed by rotating tissues by the aforementioned angle and by centering the zero coordinate to the center of the L-hBG.

Cell/tissue type enrichment analysis has been performed with EnrichR ^13^ with the ARCHS4 ^14^, CellMarkerAugmented ^15^ and Panglao ^16^ databases. These databases are incorporated within MULTILAYER, allowing to perform the aforementioned enrichment analysis directly within this tool.

### Data access

All raw datasets generated on this study have been submitted to the NCBI Gene Expression Omnibus (GEO; http://www.ncbi.nlm.nih.gov/geo/) under accession number GSE223020. Furthermore, processed spatial matrices as well as the corresponding scanned images are available herein: https://data.mendeley.com/datasets/ptv86nczbv.

## Acknowledgements

We thank all the members of our laboratories and the Genoscope sequencing platform for discussion. This work was supported by the institutional bodies CEA, CNRS, and Université d’Evry-Val d’Essonne, the Genopole Thematic Incentive Actions funding (ATIGE-2017), the ‘‘Fondation pour la Recherche Medicale’’ (FRM; funding ALZ-201912009904) as well as the Institut National du Cancer (INCa: Funding 2020-181).

## Author contributions

G Lozachmeur: formal analysis, investigation, and methodology. A Bramoulle: resources and methodology. A Aubert: resources and methodology. F Stüder: data curation, software, and formal analysis. J Moehlin: data curation, software, and formal analysis. L Madrange: resources and methodology. F Yates: writing—review and editing. JP Deslys: supervision, funding acquisition, and writing—review and editing. MA Mendoza-Parra: conceptualization, formal analysis, supervision, funding acquisition, and writing—original draft.

## Conflict of interest

The authors declare that they have no conflict of interest.

## Notes

### Competing Interest Statement

The authors have declared no competing interest.

## References

1. Rao, A., Barkley, D., França, G.S., and Yanai, I. (2021). Exploring tissue architecture using spatial transcriptomics. Nature 596, 211–220. 10.1038/s41586-021-03634-9.

2. Ståhl, P.L., Salmén, F., Vickovic, S., Lundmark, A., Navarro, J.F., Magnusson, J., Giacomello, S., Asp, M., Westholm, J.O., Huss, M., et al. (2016). Visualization and analysis of gene expression in tissue sections by spatial transcriptomics. Science 353, 78–82. 10.1126/science.aaf2403.

3. Schlecht, U., Mok, J., Dallett, C., and Berka, J. (2017). ConcatSeq: A method for increasing throughput of single molecule sequencing by concatenating short DNA fragments. Scientific Reports 7, 5252. 10.1038/s41598-017-05503-w.

4. TELP, a sensitive and versatile library construction method for next-generation sequencing - PubMed https://pubmed.ncbi.nlm.nih.gov/25223787/.

5. Shankaranarayanan, P., Mendoza-Parra, M.-A., Walia, M., Wang, L., Li, N., Trindade, L.M., and Gronemeyer, H. (2011). Single-tube linear DNA amplification (LinDA) for robust ChIP-seq. Nature Methods 8, 565–567. 10.1038/nmeth.1626.

6. Moehlin, J., Mollet, B., Colombo, B.M., and Mendoza-Parra, M.A. (2021). Inferring biologically relevant molecular tissue substructures by agglomerative clustering of digitized spatial transcriptomes with multilayer. cels 12, 694-705.e3. 10.1016/j.cels.2021.04.008.

7. Moehlin, J., Koshy, A., Stüder, F., and Mendoza-Parra, M.A. (2021). Protocol for using MULTILAYER to reveal molecular tissue substructures from digitized spatial transcriptomes. STAR Protoc 2, 100823. 10.1016/j.xpro.2021.100823.

8. Park, Y.H., Pyun, J.-M., Hodges, A., Jang, J.-W., Bice, P.J., Kim, S., Saykin, A.J., Nho, K., and AddNeuroMed consortium and the Alzheimer’s Disease Neuroimaging Initiative (2021). Dysregulated expression levels of APH1B in peripheral blood are associated with brain atrophy and amyloid-β deposition in Alzheimer’s disease. Alzheimers Res Ther 13, 183. 10.1186/s13195-021-00919-z.

9. Adnan, G., Rubikaite, A., Khan, M., Reber, M., Suetterlin, P., Hindges, R., and Drescher, U. (2020). The GTPase Arl8B Plays a Principle Role in the Positioning of Interstitial Axon Branches by Spatially Controlling Autophagosome and Lysosome Location. J Neurosci 40, 8103–8118. 10.1523/JNEUROSCI.1759-19.2020.

10. St George-Hyslop, F., Kivisild, T., and Livesey, F.J. (2022). The role of contactin-associated protein-like 2 in neurodevelopmental disease and human cerebral cortex evolution. Front Mol Neurosci 15, 1017144. 10.3389/fnmol.2022.1017144.

11. Ortiz, C., Navarro, J.F., Jurek, A., Märtin, A., Lundeberg, J., and Meletis, K. (2020). Molecular atlas of the adult mouse brain. Sci Adv 6, eabb3446. 10.1126/sciadv.abb3446.

12. Liao, Y., Smyth, G.K., and Shi, W. (2014). featureCounts: an efficient general purpose program for assigning sequence reads to genomic features. Bioinformatics 30, 923–930. 10.1093/bioinformatics/btt656.

13. Chen, E.Y., Tan, C.M., Kou, Y., Duan, Q., Wang, Z., Meirelles, G.V., Clark, N.R., and Ma’ayan, A. (2013). Enrichr: interactive and collaborative HTML5 gene list enrichment analysis tool. BMC Bioinformatics 14, 128. 10.1186/1471-2105-14-128.

14. Lachmann, A., Torre, D., Keenan, A.B., Jagodnik, K.M., Lee, H.J., Wang, L., Silverstein, M.C., and Ma’ayan, A. (2018). Massive mining of publicly available RNA-seq data from human and mouse. Nature Communications 9, 1366. 10.1038/s41467-018-03751-6.

15. Zhang, X., Lan, Y., Xu, J., Quan, F., Zhao, E., Deng, C., Luo, T., Xu, L., Liao, G., Yan, M., et al. (2019). CellMarker: a manually curated resource of cell markers in human and mouse. Nucleic Acids Research 47, D721–D728. 10.1093/nar/gky900.

16. Franzén, O., Gan, L.-M., and Björkegren, J.L.M. (2019). PanglaoDB: a web server for exploration of mouse and human single-cell RNA sequencing data. Database 2019, baz046. 10.1093/database/baz046.

