## Supplementary Figures for "A 3-dimensional molecular cartography of human cerebral organoids revealed by double-barcoded spatial transcriptomics"

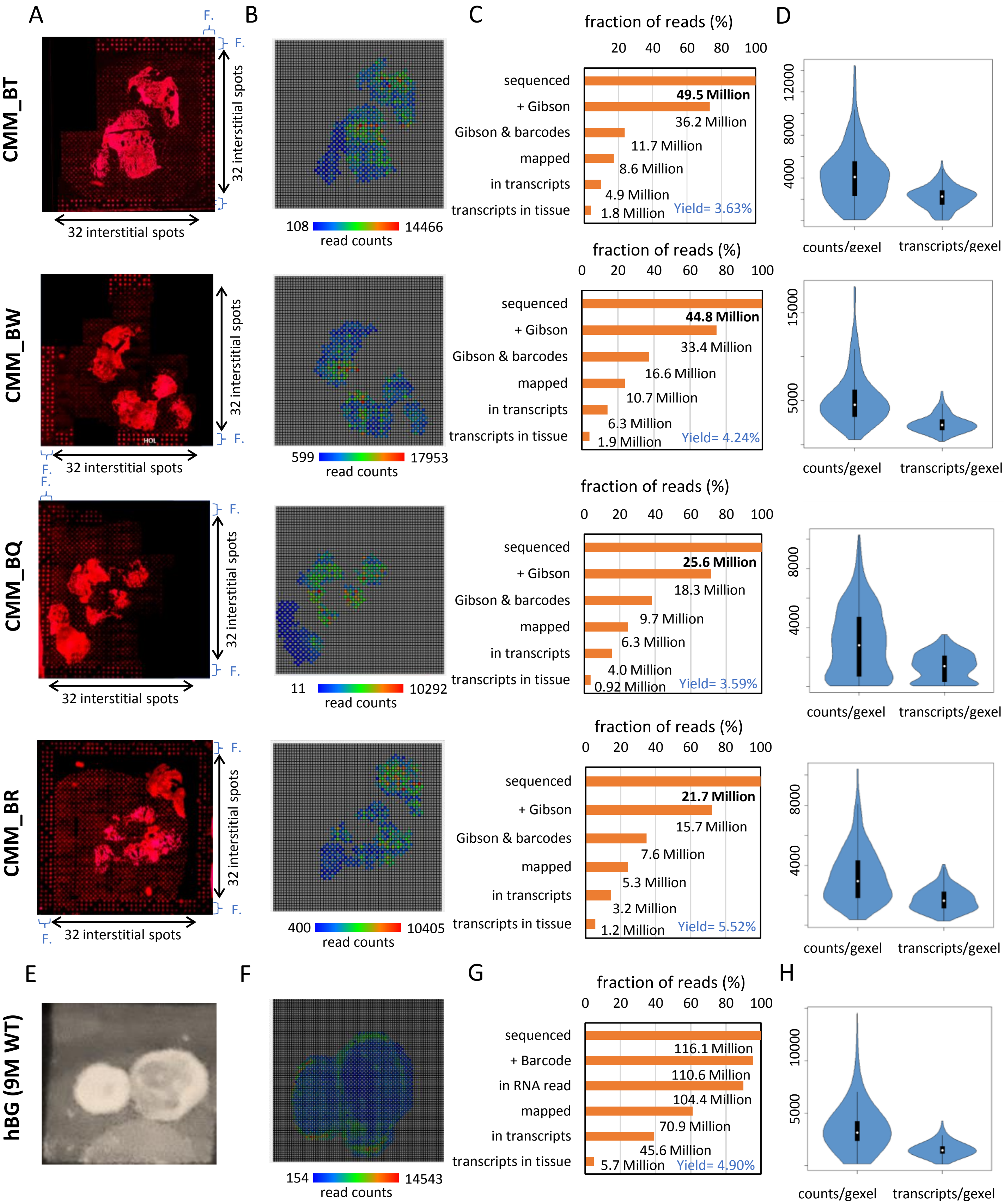


**Supplementary Figure S1: Human brain organoid tissue sections analyzed by Spatially-resolved transcriptomics in double-barcoded DNA arrays.** Four tissue sections (hBG: 4 months culture) were deposited on double-barcoded DNA arrays and sequenced at different coverage levels (21 to 49 million reads). **(A)** Micrograph of tissue sections deposited on double-barcoded DNA arrays in which cDNA synthesis is revealed with Cy3-labeled nucleotides. In addition to tissue labelling, surrounding probes can also be visualized due to the reverse transcriptase activity of adding extra Cytosine nucleotides. Fiducials corresponding to Cy3-labelled nucleotides are also visible at the borders of the micrographs (F). **(B)** Spatial transcriptomics view (MULTILAYER) composed by gexels (gene expression local pixels) associated to regions where the tissue has been localized. **(C)** Fraction of sequenced reads recovered after the primary bioinformatics treatment. **(D)** Violin plot displaying either the number of read counts or transcripts per gexel. **(E)** Bright-field micrograph corresponding to a human brain organoid (9 months) which has been analyzed with the spatial transcriptomics arrays commercialized by Visium 10x Genomics. **(F, G & H)** Similar to (B, C & D) for the Visium spatial transcriptomics readouts. Yield expressed in panel C and G corresponds to the ratio of transcripts in tissue rel. to the total sequenced reads.

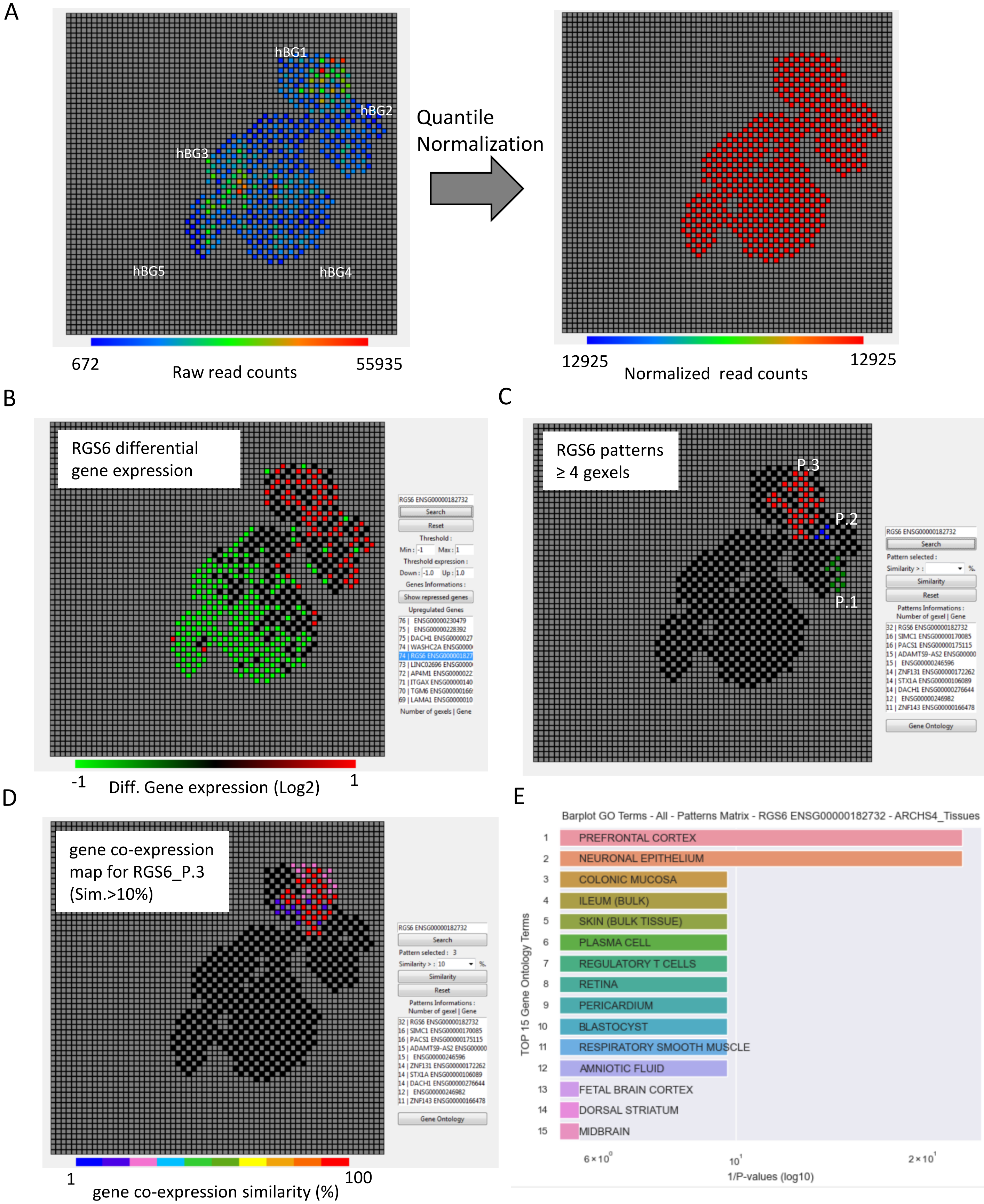


**Supplementary Figure S2: Spatially-resolved transcriptomics issued from human brain organoids analyzed with MULTILAYER (Moehlin et al; Cell Systems 2021). (A)** Left panel: Digitized view of the spatial transcriptome assessed for all four human brain organoids (hBG1 to hBG5) displaying raw read counts per gexel (local gene expression pixel). Right panel: Quantile normalization applied for correcting technical variations within raw read counts/gexel. **(B)** MULTILAYER computes differential gene expression per gexel relative to the average read counts across the tissue. MULTILAYER displays a panel (right side of the matrix) with a raking of genes based on the number of upregulated gexels (defined threshold= log2(Fold-change) >1). In (B), the differential expression levels associated to the gene RGS6 is displayed. **(C)** MULTILAYER retains gene expression patterns defined by a minimal number of contiguous up-regulated gexels (list of detected patterns displayed in the panel at the right of the matrix). Herein, MULTILAYER detected 3 patterns for RGS6. **(D)** For RGS6 pattern 3, MULTILAYER found ~30 other genes presenting upregulated patterns within the same spatial region with a co-expression similarity as low as 10% (heatmap color code associated to the displayed gexels). (E) ARCHS4 tissue terms enrichment analysis applied to co-expressed genes with RGS6 revealed signatures associated to terms like Prefrontal cortex and Neuronal epithelium, indicative of an optimal neuro-ectodermal differentiation for such region of hBG1.

**ARL8B**

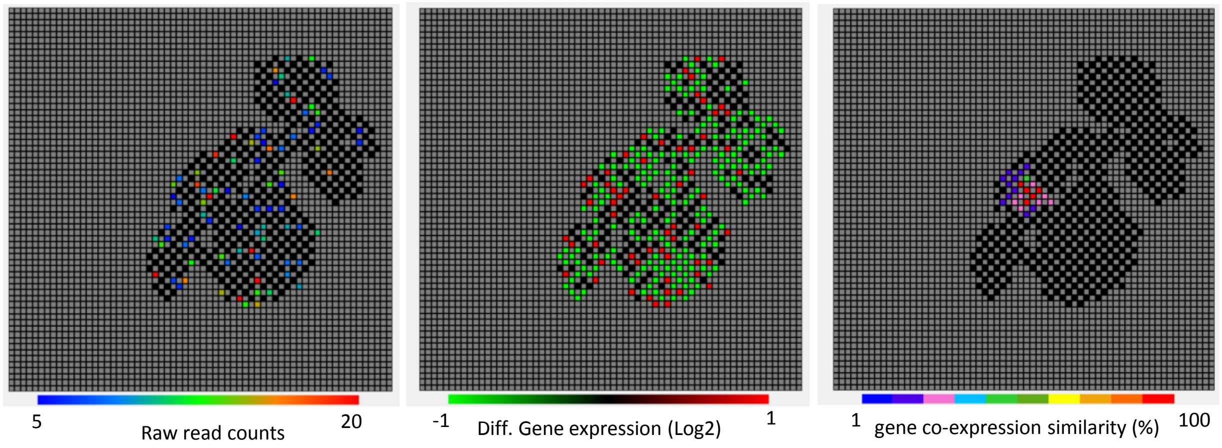


**APH1B**

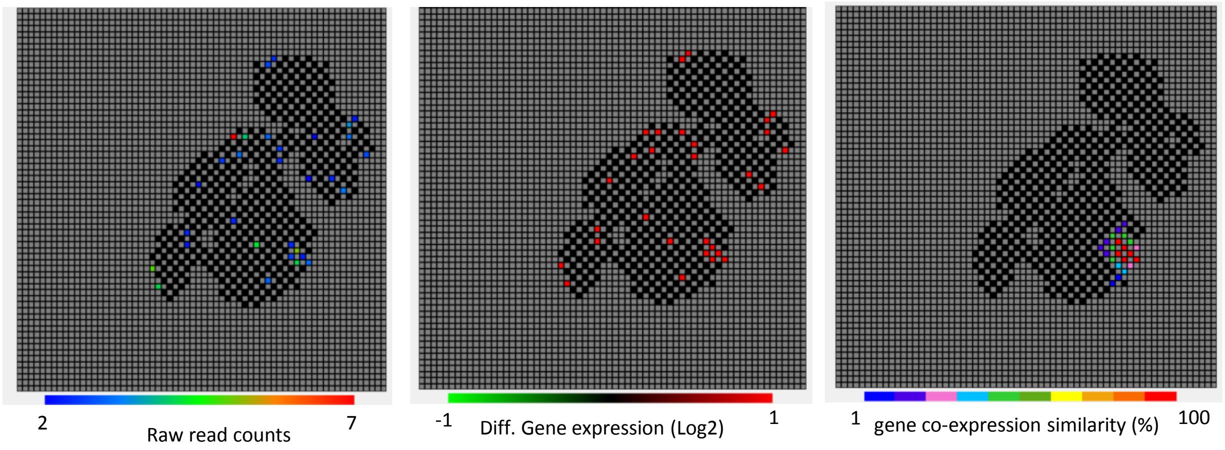


**Supplementary Figure S3: Spatially-resolved transcriptomics landscapes issued from human brain organoids analyzed with MULTILAYER (Moehlin et al; Cell Systems 2021). Top panels:** Digitized view of the spatial transcriptome map assessed for the gene ARL8B (ADP Ribosylation Factor Like GTPase 8B), known to play a key role in the positioning of Interstitial Axon Branches during the establishment of neuronal connectivity (Adnan et al; J Neurosciences 2020). Left: raw read counts per gexel (local gene expression pixel). Middle: Differential gene expression per gexel relative to the average read counts across the tissue. Right: local gene co-expression similarity (heatmap color code associated to the displayed gexels). Bottom panels: Same as top panels but for the gene APH1B which encodes for a multi-pass transmembrane protein that is a functional component of the gamma-secretase complex. Both genes do not appear up-regulated in our bulk RNA-seq assay; which might be explained by the localized induction of these genes observed in spatial transcriptomics.

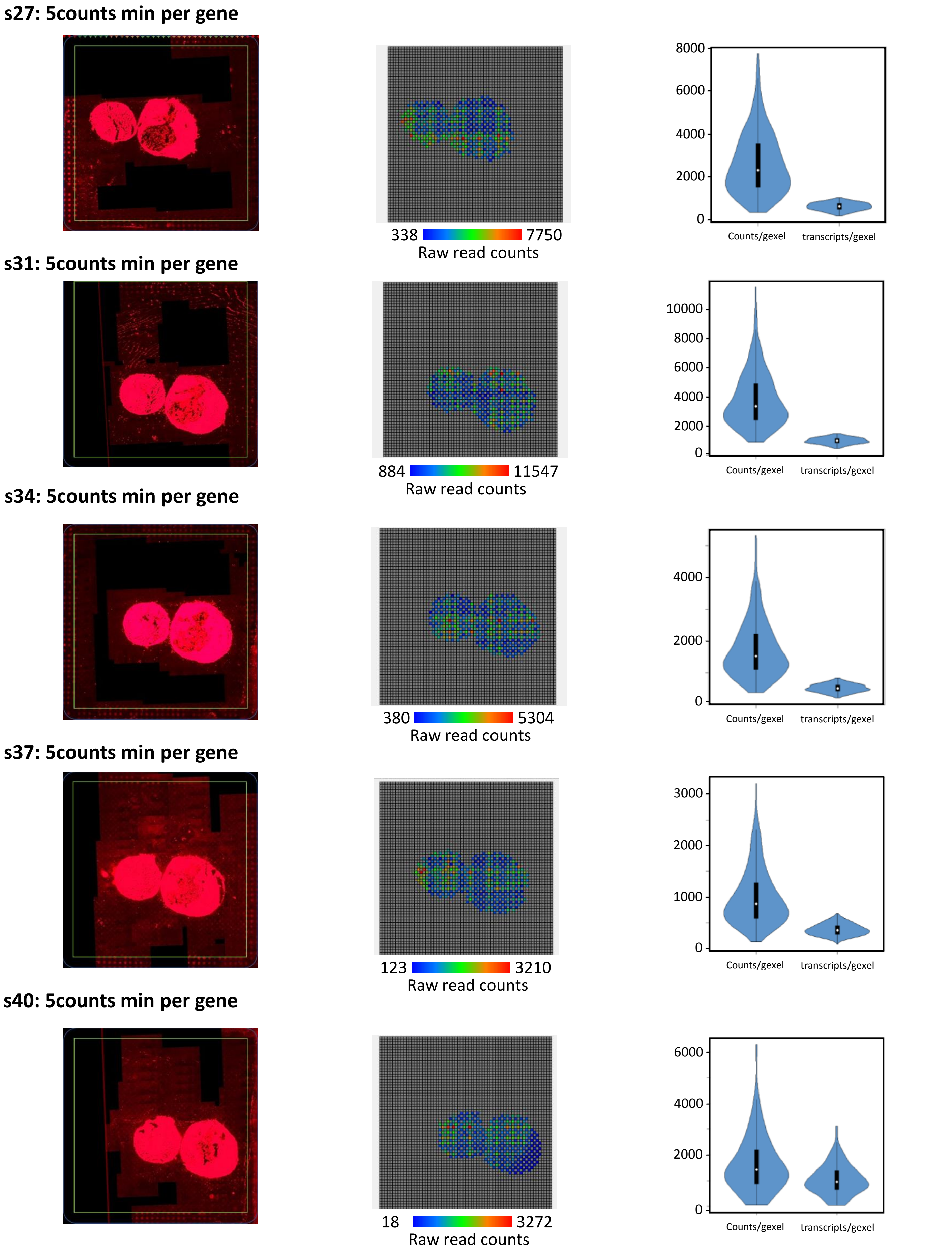


**Supplementary Figure S4: Consecutive sections analyzed by SrT for generating a 3-dimensional view of gene programs across the tissue.** Left panels: Scan of DNA arrays (TRITC filter) hosting the indicated cryosection after cDNA labelling with dCTP-Cy3. The green square defines the array region inside the fiducials. Note that not tht due to our manual scanning, not all the borders of the fiducials are available. Middle panels: SrT digitized view of the corresponding sections displaying the number of raw read counts per gexel. Right panels: Violin plots displaying either the number of read counts/gexel or the number of known transcripts/gexel.

**s44: 5counts min per gene**

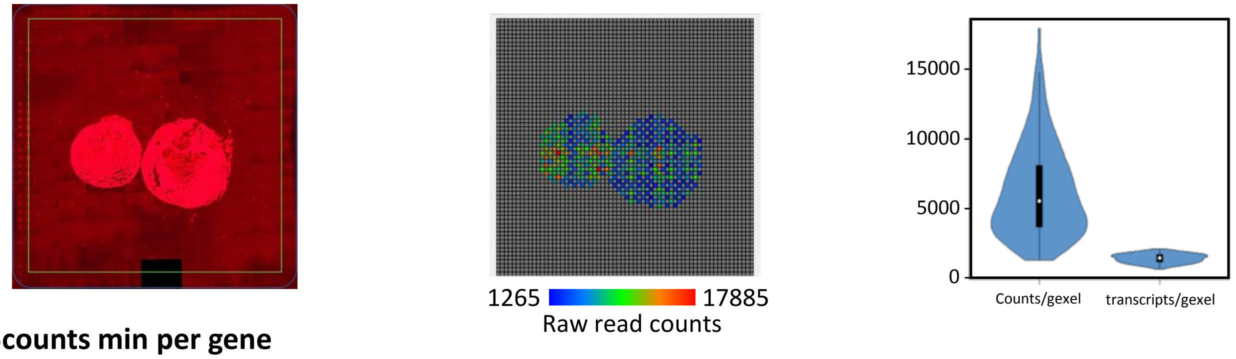


**s47: 5counts min per gene**

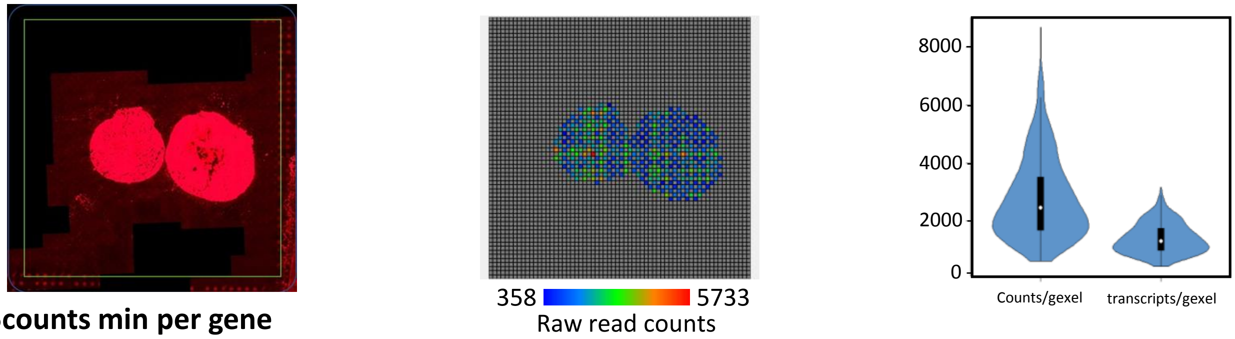


**s50: 5counts min per gene**

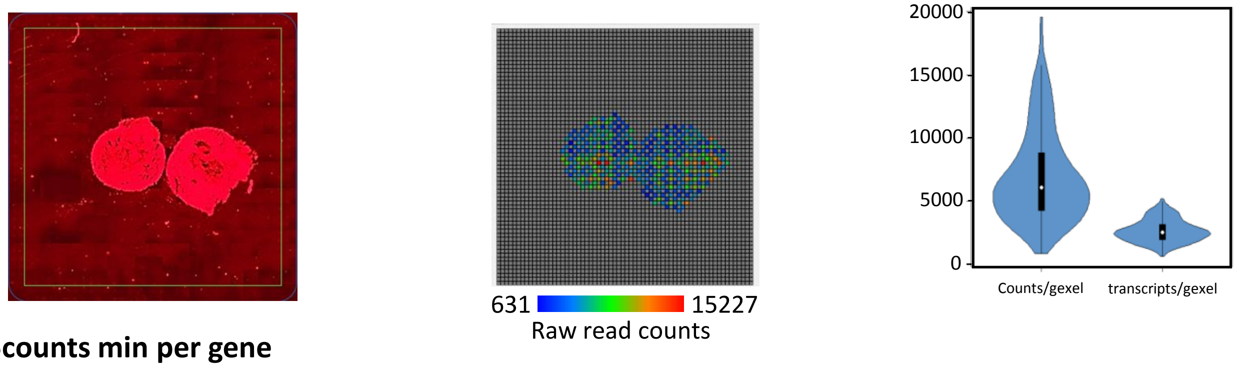


**s56: 5counts min per gene**

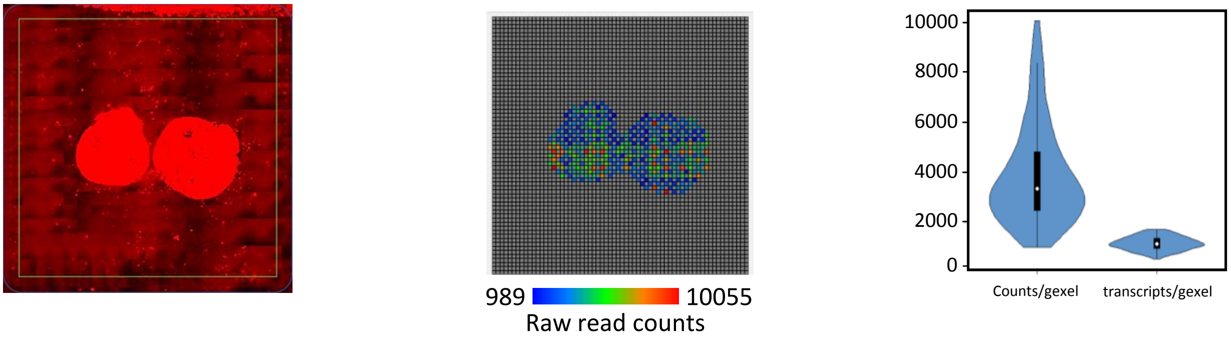


**Supplementary Figure S4: Consecutive sections analyzed by SrT for generating a 3-dimensional view of gene programs across the tissue.** Left panels: Scan of DNA arrays (TRITC filter) hosting the indicated cryosection after cDNA labelling with dCTP-Cy3. The green square defines the array region inside the fiducials. Note that not tht due to our manual scanning, not all the borders of the fiducials are available. Middle panels: SrT digitized view of the corresponding sections displaying the number of raw read counts per gexel. Right panels: Violin plots displaying either the number of read counts/gexel or the number of known transcripts/gexel.

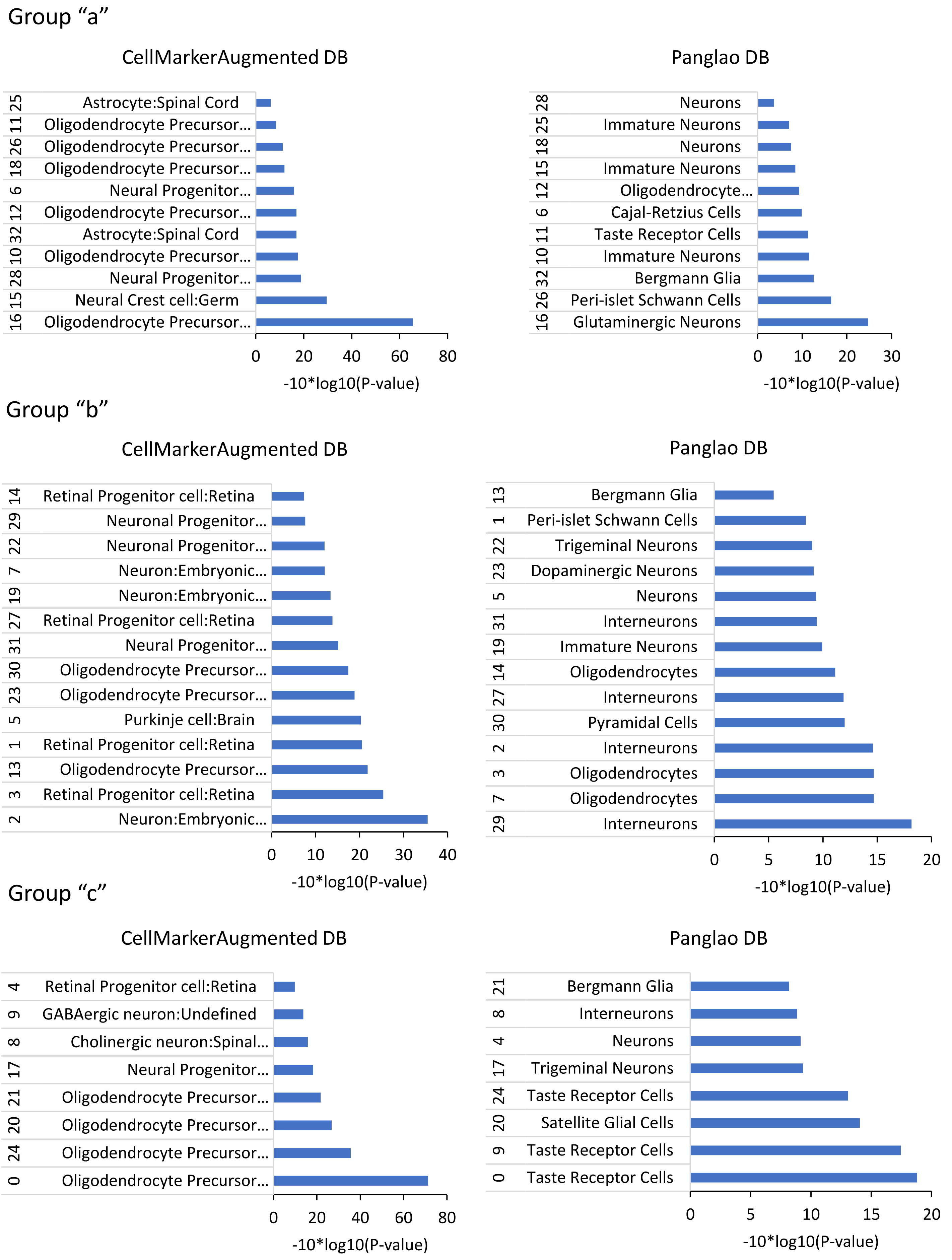


**Supplementary Figure S5: Cell/tissue types enrichment analysis performed for all three groups of community classes revealed in Figure 2.** Genes composing all 33 community classes were stratified on the grounds of their upregulation in consecutive sections. Pearson correlation clustering on the stratified genes revealed three major groups which were analyzed for cell/tissue types enrichment towards two databases: CellMarkerAugmented (Zhang et al; NAR 2019; http://xteam.xbio.top/CellMarker/) and Panglao DB( Franzen et al; Database 2019; https://panglaodb.se/). This analysis has been berformed in EnrichR where these and other GO term databases are implemented (Kuleshov et al; NAR 2016; https://maayanlab.cloud/Enrichr/).
